# The kinesin-8 Kip3 depolymerizes microtubules with a collective force-dependent mechanism

**DOI:** 10.1101/844829

**Authors:** Michael Bugiel, Mayank Chugh, Tobias Jörg Jachowski, Erik Schäffer, Anita Jannasch

## Abstract

Microtubules are highly dynamic filaments with dramatic structural rearrangements and length changes during the cell cycle. An accurate control of the microtubule length is essential for many cellular processes in particular, during cell division. Motor proteins from the kinesin-8 family depolymerize microtubules by interacting with their ends in a collective and length-dependent manner. However, it is still unclear how kinesin-8 depolymerizes microtubules. Here, we tracked the microtubule end-binding activity of yeast kinesin-8, Kip3, under varying loads and nucleotide conditions using high-precision optical tweezers. We found that single Kip3 motors spent up to 200 s at the microtubule end and were not stationary there but took several 8-nm forward and backward steps that were suppressed by loads. Interestingly, increased loads, similar to increased motor concentrations, also exponentially decreased the motors’ residence time at the microtubule end. On the microtubule lattice, loads also exponentially decreased the run length and time. However, for the same load, lattice run times were significantly longer compared to end residence times suggesting the presence of a distinct force-dependent detachment mechanism at the microtubule end. The force dependence of the end residence time enabled us to estimate what force must act on a single motor to achieve the microtubule depolymerization speed of a motor ensemble. This force is higher than the stall force of a single Kip3 motor, supporting a collective force-dependent depolymerization mechanism that unifies the so-called “bump-off” and “switching” models. Understanding the mechanics of kinesin-8’s microtubule end activity will provide important insights into cell division with implications for cancer research.

**STATEMENT OF SIGNIFICANCE:** Kinesin-8 motors are important for microtubule length regulation and are over-expressed in different types of cancer. Yet, on the molecular level, it is unclear how these motors depolymerize microtubules. Using high-precision optical tweezers, we measured how single yeast kinesin-8 motors, Kip3, interacted with the microtubule end. Interestingly, we found that single Kip3 motors were still motile at the microtubule end. The force dependence of how long single motors were associated with the microtubule end enabled us to estimate what force motors must exert onto each other to achieve the collective microtubule depolymerization speed of many motors. Our data support a collective force-dependent depolymerization mechanism. A better understanding of Kip3’s microtubule end activity has implications for cell division and associated diseases.

## INTRODUCTION

The mitotic spindle and the microtubule cytoskeleton are highly dynamic structures. During cell division, assembly and disassembly of the mitotic spindle is a crucial step. The mitotic spindle consists of cytoskeletal microtubules (MTs). Therefore, a proper control of the microtubule length is important. Microtubules are built of protofilaments—chains of tubulin dimers. These microtubules have a polar structure with a more dynamic plus end and a less dynamic minus end. There are many microtubule-associated proteins (MAPs) that are involved in MT polymerization (1), stabilization (2, 3), and depolymerization (4–6) and, thus, MT length control. Furthermore, members of the kinesin-8 family are involved in control of the MT architecture (7–10), knock-out mutants in budding yeast exhibit unnaturally long microtubules (6), and an overexpression in humans associated with cancer (11, 12). While high concentrations of the budding yeast kinesin-8 *Kip3* lead to collective, length-dependent microtubule depolymerization (5, 6, 13), individual motors at low Kip3 concentrations have been proposed to stabilize MT plus ends (14). The length-dependent depolymerization is explained by an antenna model and results from the combination of the microtubule depolymerase activity with the very high processivity towards the microtubule plus end and average run length of 12 µm (5, 15). Compared to conventional kinesin-1, Kip3 is slow and stalls already at a force of ≈1.2–1.5 pN (16, 17). Kip3 also has an additional MT-binding site at its tail that enables it to cross-link MTs (15) and together with its weakly-bound slip state (16) contributes to its extraordinarily long run length. However, how kinesin-8 depolymerizes microtubules is still not clear and under debate.

In one model, Varga et al. suggested that Kip3 depolymerizes microtubules with a *bump-off* mechanism (13) in which one Kip3 is waiting at the MT end until another Kip3 reaches the microtubule end and bumps the waiting Kip3 off the microtubule. The model suggests that the bumped-off Kip3 takes one to two tubulin dimers with it. This model is motivated by the observation that the long microtubule end residence time of single Kip3 decreases exponentially with higher Kip3 concentrations, accompanied by increased MT depolymerization. The bump-off model requires Kip3 to reach the MT end. While this feature can be well explained by Kip3’s long run length and its suggested ability to bypass obstacles on the microtubule (17–20), the activity of Kip3 at the MT end, i.e. the depolymerization mechanism itself, is poorly understood.

In contrast, a more recent study by Arellano et al. proposed a different *switching* model (21). The authors measured that Kip3 has a higher affinity for curved tubulin protofilaments. In addition, once Kip3 is bound to curved tubulin dimers, the motor’s ATPase activity is suppressed. Therefore, Kip3 accumulates at MT ends where single protofilament overhangs are curved and unstable and, thereby, promotes MT depolymerization through strong binding to curved protofilaments. This model is supported by the finding that monomeric, immotile Kip3 can promote MT depolymerization by binding directly to the microtubule end showing that motility is not required to reduce end residence times. However, the switching model does not rule out a cooperative bump-off model since high motor concentrations were used and motors, instead of walking to the last bound Kip3, may have bound directly next to the last motor out of solution. The observed switch of Kip3 into a non-motile state at the microtubule end raises the question how Kip3’s binding state at the microtubule end differs from the state on the microtubule lattice under different nucleotide conditions. Moreover, the influence of loads or pushing forces on Kip3 at the microtubule end—that are expected to be present if large numbers of Kip3 push each other from the microtubule in a kind of traffic jam—is unknown.

Here, we used high-precision optical tweezers (22) in a two-dimensional force-feedback mode (23) to apply constant assisting and hindering loads along the microtubule axis on motile Kip3 while stepping towards and remaining at the microtubule end. We found that both the end residence time and the forward/backward motion at the microtubule end decreased exponentially with higher loads. Controls with Kip3 walking on the microtubule lattice and bound to microtubules in the presence of ADP and AMPPNP, respectively, showed that the detachment mechanism from the microtubule end is distinct. Altogether, we propose that Kip3 depolymerizes microtubules with a collective, force-dependent mechanism requiring piconewton loads to effectively remove a tubulin dimer. This motor-driven depolymerization mechanism combines and necessitates both the bump-off and the switching models.

## MATERIALS AND METHODS

### Microtubule preparation

Porcine tubulin (3 µM) was polymerized in PEM buffer (80mM PIPES, 1mM EGTA, 1mM MgCl_2_, pH = 6.9) with 1mM MgCl_2_ and 1mM guanosine-5’-[(*α,β*)-methyleno] triphosphate (GMPCPP, Jena Bioscience, Jena, Germany) for 1.5 h at 37 °C. Afterwards, the GMPCPP-stabilized microtubules were diluted with PEM, spun down in a Beckman airfuge, and resuspended in PEM. If not noted otherwise, all chemicals were from Sigma. Microtubules were visualized with differential interference contrast employing a light emitting diode (LED-DIC) (24). For total internal reflection fluorescence (TIRF) assays, 10 % rhodamine-labeled tubulin was used to image microtubule growth.

### Functionalized microsphere preparation

To specifically bind enhanced green fluorescent protein (eGFP)-tagged proteins, carboxylated polystyrene microspheres (mean diameter 0.59 µm, Bangs Lab., Fishers, USA) were coated covalently with a small fraction of anti-GFP nanobody (kindly provided by Ulrich Rothbauer, NMI, Reutlingen, Germany), surrounded by monofunctional PEG molecules (mPEG, 2 kDa, Rapp Polymere, Tübingen, Germany) in a ratio of 1: 10^4^, based on protocols described in (25) and as depicted in Fig. 1A. The mPEG forms a polymeric brush that suppresses unspecific binding.

**FIGURE 1.**
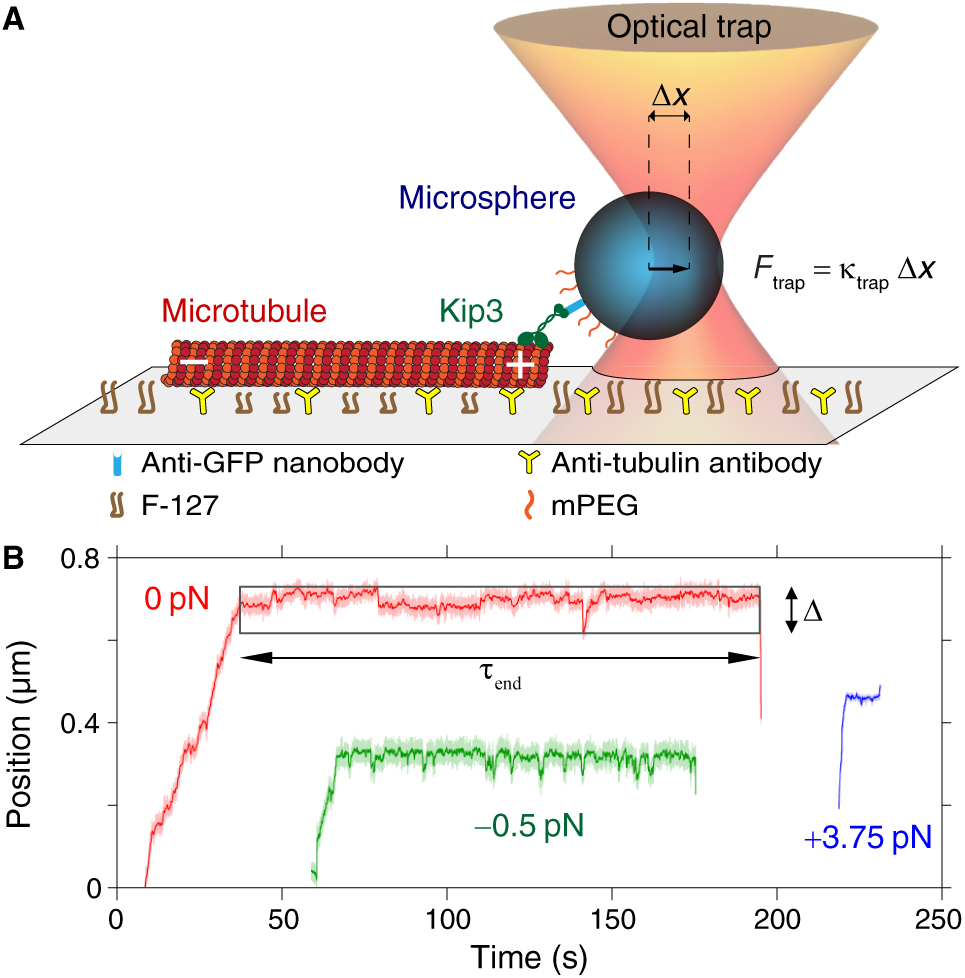
Probing the activity of Kip3 at the microtubule plus end using optical tweezers. (*A*) Schematic of the Kip3 stepping assay (side view, not to scale). Loads *F* were applied along the MT axis until Kip3 detached from the microtubule end. (*B*) Exemplary traces of Kip3 on the microtubule for three different loads offset for clarity (assisting and hindering loads are positive (blue) and negative (green), respectively). Motors walked to the microtubule plus end, stepped forward and backward after reaching it, and finally detached from the MT end. The end residence time *τ*_end_ was calculated as the time between the arrival at and detachment from the microtubule end. The end range ∆ was calculated as the difference between maximal and minimal position while the Kip3 was at the microtubule end.

### Sample preparation and assay

Experiments were performed in flow cells that were constructed using silanized, hydrophobic glass cover slips and parafilm as described before (19). Full-length budding yeast Kip3 (Kip3-eGFP-His_6_) was expressed and purified according to (16, 26). The motility buffer for Kip3 stepping assays was PEM supplemented with 1mM ATP, 0.1 mg/ml casein, 112.5mM KCl, and an anti-fading mix (10mM DTT, 20mM glucose, 20 µg/ml glucose oxidase, 8 µg/ml catalase) (16, 19, 25). Experiments with motility buffer and different nucleotides apart from ATP used the same nucleotide concentration (1 mM) unless mentioned otherwise. Functionalized microspheres were mixed with motor proteins in motility buffer to a motor-to-microsphere ratio for which every third microsphere showed motility, implying single-molecule conditions with 95 % confidence (25, 27). The channels of flow cells were washed with PEM, filled and incubated successively with anti *β*-tubulin I (monoclonal antibody SAP.4G5 from Sigma in PEM), Pluronic F-127 (1 % in PEM), and microtubules in PEM. Finally, the kinesin-microsphere suspension was flowed in.

### Optical tweezers setup

Measurements were performed in a single-beam optical tweezers setup combined with LED-DIC to visualize single microtubules as described in (22, 24, 28). The setup is equipped with a millikelvin precision temperature control (22) set to 29.500 °C and a two-dimensional (2D) force clamp with a feedback rate of 1 kHz and a sampling rate of 4 kHz as described in detail in (23). Forces along the microtubule axis, are denoted by *F*. The optical trap was calibrated by a combined power spectral density–drag force method (29, 30). We used a lateral trap stiffness of 0.03–0.05 pN/nm with a laser power in the trapping focus of about 20–30 mW.

### Tracking the activity of Kip3 at the microtubule plus end and on the microtubule lattice

Kip3 experiments were performed in the way described previously (16, 17, 19). Briefly, Kip3-coated microspheres were trapped and positioned on top of microtubules to await Kip3-initiated motility. When the motor started to pull on the microsphere, the 2D force clamp was switched on with assisting or hindering loads applied on the kinesin along the MT axis, denoted as positive and negative, respectively, as illustrated in Fig. 1A. The 2D feedback ensured that no sideward loads were applied. Kip3’s zero-load speed of ≈55 nm/s and stall force of about 1.5 pN were consistent with previous results (16, 17, 19). Microsphere positions were processed with a running median filter over 500 data points. We recorded a total of 510 traces of 93 different single-Kip3-powered microspheres. We defined the arrival of Kip3 at the microtubule plus end as the time point when apparent directed microsphere motion stopped. By visual inspection of the traces and LED-DIC images, it was confirmed that indeed the microtubule end was reached. The time between arrival and detachment from the microtubule end was defined as end residence time *τ*_end_ (Fig. 1B). During this period, we determined the end range ∆ by first subtracting the minimal from the maximal filtered position. To correct this difference for non motility-related, load-dependent Brownian noise in the position, we measured this noise of immotile Kip3-coated microsphere positions on microtubules and subtracted the peak-to-peak noise, calculated as six standard deviations, from the latter difference resulting in the end range ∆. Note that hindering loads of less than 1pN were not used for microtubule end tracking experiments because such loads are close to the stall force and Kip3 rarely if at all reached the MT plus end under these conditions. As a precise measure for the size of steps that occurred in the end range including multiples of the underlying, unknown step size *δ*_0_, we used the function *µ*^2^(*δ*) on the step size data *δ*. The step size *δ* was determined from peak distances of a multi-Gaussian fit to position histograms. We calculated the measure according to 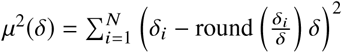, where the round function rounds the argument to the nearest integer value. *µ*^2^(*δ*) has a minimum at the unknown value of the step size *δ*_0_.

To measure microtubule depolymerization speeds, we repeatedly performed Kip3 runs to the microtubule end in trapping assays using the same microsphere on the same microtubule with the same starting position and continuous data acquisition for up to 10 min. Each run lasted on average about 1 min. For each run, the maximal position was considered as the position of the MT plus end. The slope of lines fitted to these MT end positions versus their corresponding time points resulted in the depolymerization speed. According to the defined geometry, depolymerization speeds were denoted as negative. Since depolymerization speeds did not signficantly depend on load and direction, we pooled all depolymerization speed data. As a control on the MT lattice, we measured the run time and run length of Kip3 as a function of assisting loads (243 runs with 19 microspheres). Since the mean run length of Kip3 at zero load exceeded the range of our force-clamp, we were only able to measure run times and lengths for loads ≥2.5 pN. For these loads, run lengths were sufficiently small. Included in the MT lattice analysis were runs that had slip events (16); excluded in the lattice analysis were runs in which Kip3 reached the MT end or walked out of the force-clamp range. In another control, we measured the attachment time of Kip3 (123 events of 17 microspheres) as a function of load in the presence of the non-hydrolyzable ATP analog adenosine-5’-[(*β, γ*)-imido] triphosphate (AMPPNP, Jena Bioscience). With AMPPNP, kinesins bind tightly to the MT lattice (31). For this assay, the motility buffer contained 1mM AMPPNP instead of ATP. Because the attachment times in the presence of AMPPNP were very long, they could only be measured for loads ≥3pN. Even for these loads, a few events exceeded the maximal measurement time or were initiated at a lower load and then due to the lack of detachment the load was increased. Using this final load or maximal measurement time, these events were included in the analysis. Thus, measured times may be an underestimation of the true attachment times. Without kinesin motility in the presence of AMPPNP, the polarity of the used microtubules was unknown. Therefore, loads were applied to Kip3 on the same microtubule repeatedly in both directions along the MT axis until the Kip3-coated microsphere detached. For a load of 4.25 pN, we did not find a significant asymmetry between the attachment times as a function of load directions on a single microtubule (Mann-Whitney U-test, *p* = 0.36, number of attachment times for each direction *N* = 15). Thus, attachment times for both load directions were pooled and analyzed together. Data for ADP were also pooled (32).

### TIRF stepping assays as control

As a control for Kip3 activity in the absence of load, we measured the end residence time, run time and run length of single GFP-tagged Kip3 in a TIRF microscope on rhodamine-labeled microtubules as described in (25). The mean end residence time was 80 ± 10 s (standard error if not noted otherwise, *N* = 58), the mean run time was 150±30 s (*N* = 26), and the mean run length was 12 ± 1 µm (*N* = 47). The end residence time, run time, and run length were consistent with literature values (5, 13, 15). Because the GFP nanobody has a *K*_*d*_ of about 0.6 nM (33), we expect that in trapping assays under thermal equilibrium and single-molecule conditions some of the Kip3 motors are not bound to the GFP-nanobody– functionalized microspheres, but are free in solution. These free motors are not seen in the optical trapping assay but may contribute to MT depolymerization. Therefore, we used the Kip3-microsphere motility suspension and measured the Kip3 flux to the microtubule plus end in a TIRF microscope. By counting the number of Kip3 run events to the microtubule plus end per time and microtubule, we measured a flux of about 0.018 Kip3/min/MT. Thus, only about once per hour a free Kip3 motor might contribute to MT depolymerization in the optical tweezers assay. Therefore, we can rule out a significant effect of free motors during trapping assays.

### Spontaneous microtubule depolymerization

Spontaneous depolymerization of the microtubule plus end of immobilized, GMPCPP-stabilized microtubules was measured by image analysis of long-term videos recorded with LED-DIC microscopy over >1 h. In control measurements, the trapping laser was turned on and positioned over the microtubule end. The laser power was about 30 mW in the laser focus. No microsphere was trapped and Kip3 was not in the motility buffer. Images were analyzed with Fiji (34).

## RESULTS

### Kip3 was motile at the microtubule end taking 8-nm forward and backward steps

To determine the activity of Kip3 at the microtubule end, we tracked microspheres functionalized with single motors as a function of time while applying constant loads with the optical tweezers (Fig. 1A). To this end, we first placed motor-coated microspheres several micrometers away from the end of the microtubules and engaged a 2D force clamp (23). Conveniently, due to the high processivity of Kip3, motors reliably walked to and localized at the MT plus end with nanometer precision. Surprisingly, once at the microtubule end, microspheres did not stop but continued a slow forward and backward movement before finally detaching from the microtubule (Fig. 1B). Apart from this slow motion and also during the approach to the microtubule end, we observed motor slipping in the direction of loads consistent with previous results (16). During the end residence time, we determined the difference between maximal and minimal mean position that we call the end range ∆ (Fig. 1B). In the absence of loads and for hindering loads, this distance was on the order of 100 nm and showed a large variation that we attribute to slip events (Fig. 2A). With assisting loads, the end range decreased and reached a constant value of 7.8±0.8 nm (*N* = 6) suggesting discrete, single 8-nm, center-of-mass steps typical for kinesin motors. Indeed, with a magnified view of the end motion, discrete steps are visible (Fig. 2B,C). Based on a histogram analysis, the motion—including normal slow steps and fast slipping events—occurred in multiples of about 8 nm (Fig. 2D–F). While there was forward and backward motion at the microtubule end, there was no net bias to the motion. The mean velocity at the microtubule end was not significantly different from zero (confirmed by a *χ*^2^-test, 0.2 ± 0.1 nm/s, *N* = 15, Fig. S1). Therefore, microtubules were on average neither growing nor shrinking during the residence time. Together, before detachment, Kip3 was still motile at the MT end taking several 8-nm forward and backward steps with no net motion.

**FIGURE 2.**
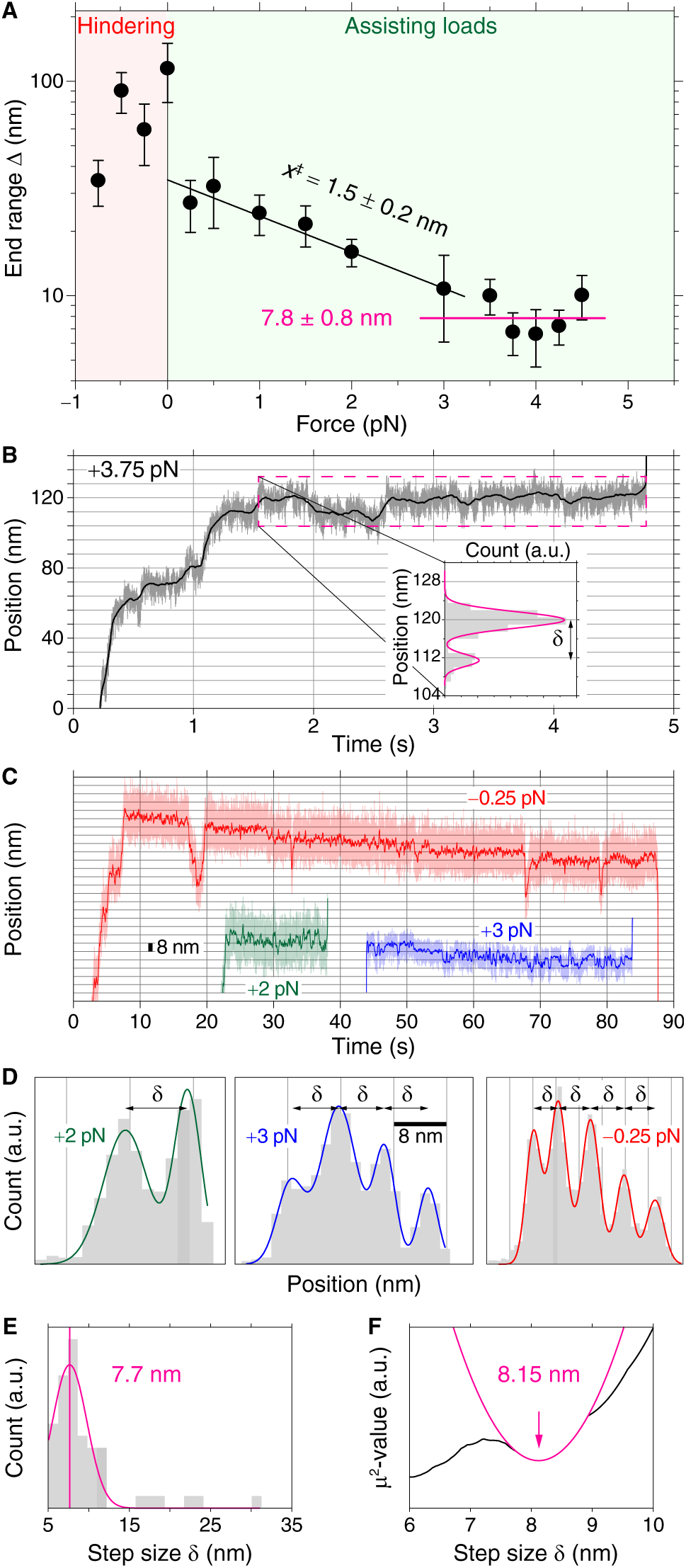
End range of stepwise forward/backward motion of Kip3 at the MT plus end decreased with load. (*A*) End range ∆ (full black circles) as a function of load with exponential fit (black line). Error bars are standard errors. The horizontal magenta line refers to a weighted mean of end ranges for loads >3 pN with weighted mean and standard error given. (*B,C*) Exemplary traces for indicated loads of Kip3 stepping to the microtubule plus end showing forward and backward steps during the end residence. Inset: Histogram of filtered position at the microtubule end with multiple Gaussian fits (magenta line). The step size *δ* is defined as difference between the peak positions of the Gaussians. (*D*) Histograms of Kip3 positions for corresponding traces in *C* with multiple Gaussian fits on 8-nm grids. (*E*) Histogram of all step sizes *δ* with a Gaussian fit and peak position at 7.7 ± 0.4 nm (*N* = 54 from 33 traces, *magenta line*). (*F*) Measure *µ*^2^ as a function of possible step sizes *δ* (*black line*) with a local minimum fitted by a parabola (*magenta line*) with a center at 8.15 ± 0.01 nm.

### The end residence time of Kip3 exponentially decreased with assisting loads

Since the end residence time of free, single Kip3 motors decreased exponentially with increasing motor concentration (13), we asked the question whether this crowding effect is due to forces that motors exert onto each other as suggested by the bump-off model and whether we can achieve the same effect by applying an external load on a single Kip3 at the microtubule end. Thus, we measured the end residence time as function of load. For all loads, the end residence time was exponentially distributed (see Fig. S2 for examples). The mean end residence time of 64 ± 9 s for Kip3 without load was consistent with the TIRF microscopy control and literature values (13–15). Load decreased end residence times (Fig. 3A, black circles). Intriguingly, only for assisting loads larger than about 0.5 pN, end residence times decreased exponentially. Based on fits using the Bell model 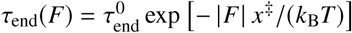, we determined the distance to the transition state *x*^‡^ for assisting and hindering loads to be 2.9 ± 0.1 nm and = 4.3 ± 0.6 nm, respectively, where *k*_B_ is the Boltzmann constant and *T* is the absolute temperature. These values differed significantly (*p* = 0.006) suggesting a bias on load direction. All measured transition distances *x*^‡^ are indicated in Fig. 2 and Fig. 3 and are summarized in Table S1. To test whether sideward loads also reduced the end residence time, we analyzed data from previous side-stepping assays (19). Without any load applied along the MT axis, but with a sideward load of 0.5 pN, we measured an end residence time of 33 ± 3 s (SEM, *N* = 49, 30 microspheres, open black circle in Fig. 3A). This time is significantly smaller than the unloaded case (confirmed by a Mann-Whitney U-test, *p* = 0.03), comparable to that of hindering loads that have the same magnitude, but larger compared to the equivalent assisting loads. Thus, force acting in the direction of microtubule growth had the largest effect. Together, the load dependence of the end residence time suggests that motor detachment from the end is susceptible to force and/or that removal of tubulin dimers is facilitated by force.

**FIGURE 3.**
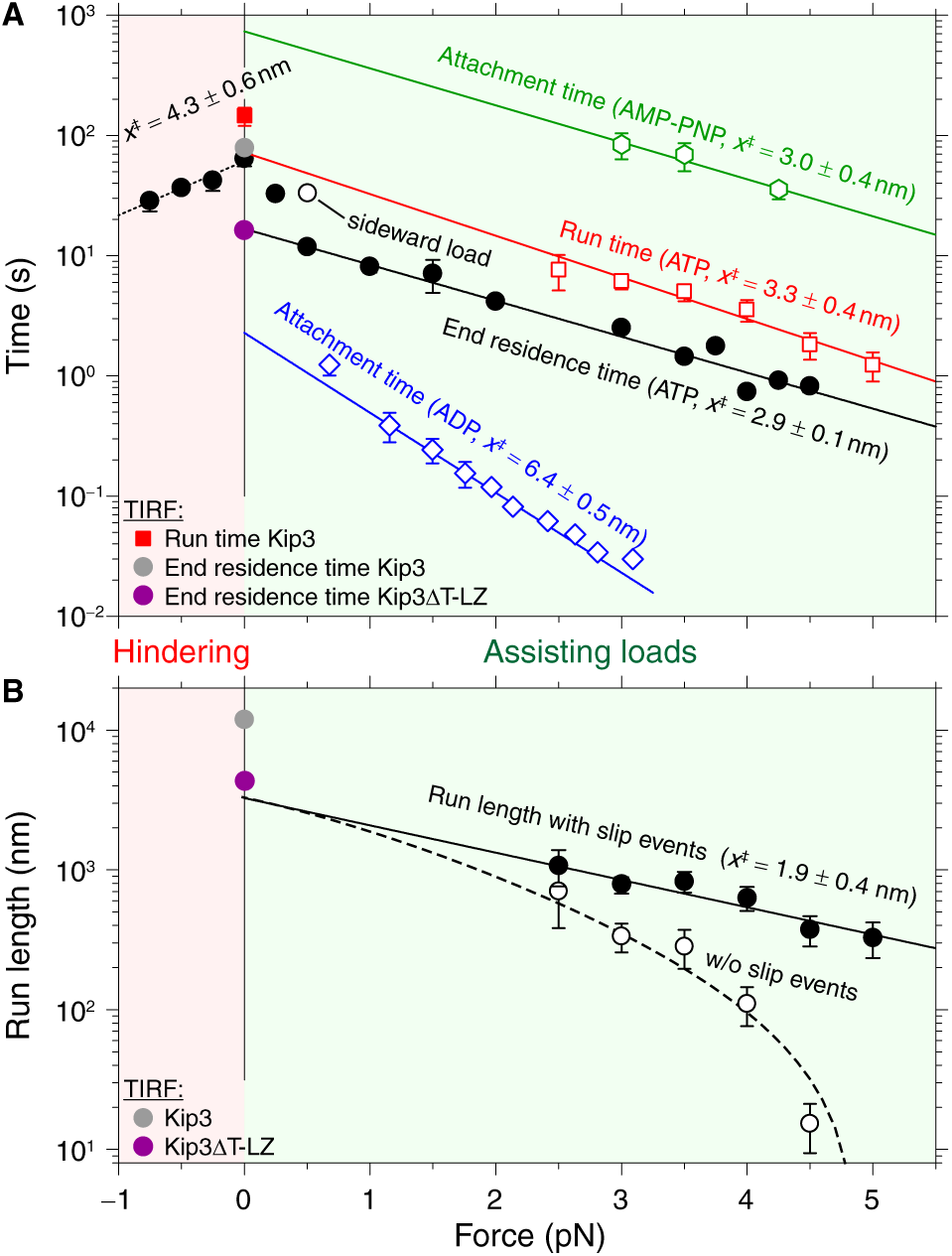
Microtubule end residence time of Kip3. (*A*) End residence time in ATP (*black circles*) with exponential fits for hindering loads (*black dotted line*) and assisting loads >0.5 pN (*black line*), run time on the MT lattice in ATP (*open red squares*) with exponential fit (*red line*), attachment time in AMPPNP (*open green hexagons*) with exponential fit (*green line*), attachment time in ADP (*open blue diamonds*, (32)) with exponential fit (*blue line*), and end residence time under sideward loads (*open black circle*, (19)). In the absence of load, mean end residence time (*grey circle*) and run time (*red square*) from TIRF measurements together with the literature value for the end residence time of truncated Kip3∆T-LZ (*purple circle*, (14)) are plotted. (*B*) Run length (*black circles*) of Kip3 as a function of load with exponential fit (*black line*) together with the zero-force run length measured in TIRF assays (*grey circle*) and run lengths from the literature (14) for truncated Kip3∆T-LZ (*purple circle*). From the original run length data that included slipping events, the average cumulative slip distance from (16) was subtracted and plotted as *open black circles* together with a model (*black dashed line*, see text). All error bars are SEMs.

### Kip3 detachment from the MT end was different from MT lattice

To test whether force-induced detachment is faster at the microtubule end compared to the lattice, we investigated how the attachment time of Kip3 at the microtubule end differed from the one on the lattice. Therefore, we measured the run time in the presence of ATP as a function of assisting load on the microtubule lattice (open red squares in Fig. 3A). We found that also the mean run times on the MT lattice decreased exponentially with increasing assisting load. In this case, the distance to the transition state was 3.3 ± 0.4 nm comparable to that of the end residence time. However, at the same load, the run times themselves were significantly longer compared to the end residence times. For example, at an assisting, MT plus-end directed force of +4.5 pN, the lattice run time was more than a factor of two longer compared to the end residence time suggesting that the motor detachment mechanism from the lattice differed from the one at the microtubule end. Interestingly, the run times on the lattice extrapolated roughly to the zero-force end residence time and run time on the lattice. To compare end residence times in ATP to that of motors in a strongly bound state, we measured the attachment time of Kip3 on the MT lattice in the presence of AMPPNP—a non-hydrolyzable nucleotide analog of ATP— as a function of load (open green hexagons in Fig. 3A). The exponential force dependence resulted in a distance to the transition state of 3.0 ± 0.4 nm again similar to the other cases above. More importantly, the AMPPNP attachment times were more than an order of magnitude longer compared to the run times and end residence times in ATP. Finally, to show that Kip3 does not slip from the MT end in the weakly-bound ADP state, we compared the end residence times with the attachment times of Kip3 on the MT lattice in the presence of ADP (open blue diamonds in Fig. 3A). For this purpose, we reanalyzed data from Bormuth *et al.* (32), where Kip3 was pulled over microtubules in the presence of 2mM ADP. Also, these attachment times decreased exponentially under loads. For ADP, the distance to the transition state of 6.4 ± 0.5 nm was significantly larger compared to the other cases. Strikingly, the attachment times were at least an order of magnitude shorter than the end residence time. However, even if Kip3 slipped in a weakly-bound state to the microtubule end, motors remained there without instantaneous detachment (Fig. S3). Together, the data in different nucleotide states suggest that the diminished Kip3 end residence time is not due to sliding off the microtubule end, but due to force that is acting on the motor.

### Repeated microtubule end tracking on the same microtubule is consistent with motor-induced tubulin removal

To test whether we could measure depolymerization due to single-motor microtubule end activity, we repeatedly tracked the same Kip3-coated microsphere on the same microtubule (Fig. 4A). We observed that the maximal position along the microtubule—i.e. the position of the MT end—decreased over time. Based on a linear fit to the maximal positions over time, we determined the depolymerization speed. Averaged over both hindering and assisting loads, the mean depolymerization speed was −0.33 ± 0.03 nm/s (*N* = 63 on 31 microtubules, Fig. 4B). By tracking microtubule depolymerization using LED-DIC (24) under identical conditions, but in the absence of Kip3 with and without trapping laser, the spontaneous, total depolymerization speed including depolymerization from the minus and plus end was −0.20 ± 0.01 nm/s (*N* = 137, Fig. 4B). The depolymerization speed was not significantly affected by the trapping laser at the used power. The spontaneous depolymerization speed was comparable to the literature value of ≈ −0.25 nm/s (13), but significantly smaller compared to the mean value obtained with single Kip3 motors under load (*t*-test, *p* < 10^−6^). Thus, single Kip3 motors subjected to loads increased the microtubule depolymerization rate suggesting that end detachment promotes depolymerization.

**FIGURE 4.**
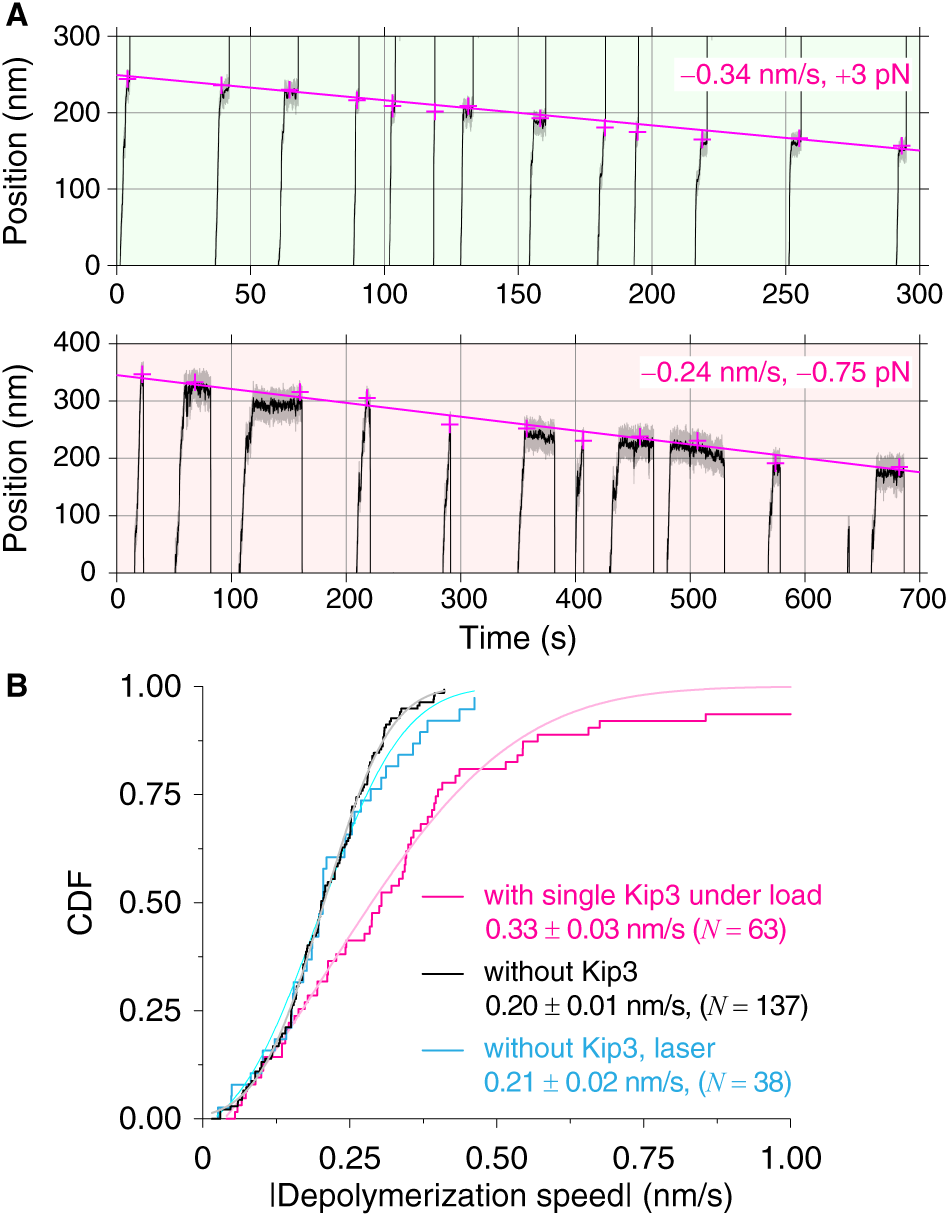
Motor-induced and spontaneous microtubule depolymerization. (*A*) Two exemplary position measurements of multiple runs of Kip3 to the MT end each starting from the same position on the same microtubule under loads of +3 pN (assisting, top) and −0.75 pN (hindering, bottom), respectively. For each run, the maximal position and corresponding time point (*magenta crosses*) were determined as a measure for the MT end position as a function of time. Linear fits of these data points (*magenta lines*) were used to calculate the MT depolymerization speeds. (*B*) Cumulative distribution function (CDF) of the absolute value of MT depolymerization speed and CDF normal distribution fits (as guide to the eye) to the data. Plus end MT depolymerization with single Kip3 under load (*magenta lines*). Total MT depolymerization (plus and minus end together) without Kip3, without and with trapping laser on (*black* and *blue lines*, respectively). Means, standard errors of the means, and numbers of data points are given.

### The run length of Kip3 decreased exponentially with load

To gain further insight into the detachment process, we also measured the load-dependent Kip3 run length on the MT lattice (Fig. 3B). We found that also the mean run length decreased exponentially with assisting loads. The distance to the transition state was 1.9 ± 0.4 nm, smaller compared to all other transition distances. The extrapolated zero-load run length of 3.3 ± 1.0 µm was significantly shorter compared to the directly measured one (TIRF microscopy and literature values on the order of 10 µm) but comparable to the value of 4.3 ± 0.2 µm reported for a Kip3 mutant, Kip3∆T-LZ, that lacks the tail microtubule binding site (14). During run events, we observed slip events consistent with previous work (16). Even though slips last only for a few milliseconds (16), the slip distances accumulate to a significant amount during a run. We estimated the mean cumulative slip distance based on a multiplication of the force-dependent slip distance and frequency (16) with the fitted run time (solid line in Fig. 3B). When we subtracted this cumulative slip distance from the mean run length, the slip-corrected run length (dashed line in Fig. 3B) was more sensitive to force confirming the notion that the slip state acts as a molecular safety leash (16). The strong force dependence also suggests that motors on the lattice subjected to high assisting loads mostly translocate in the weakly-bound slip state and lattice runs are consequently also terminated via this slip state in contrast to the observation that motors did not slip off the microtubule end (Fig. S3).

## DISCUSSION AND CONCLUSION

At microtubule plus ends, we tracked the activity of single Kip3 motors with high precision under varying loads. Interestingly, Kip3 was still motile at the microtubule end taking several 8 nm forward and backward steps before detachment. The distance that the motor explored at the microtubule end decreased with applied loads, whereas the reduction was larger for assisting loads compared to hindering loads. For hindering loads exceeding the stall force, motors did not reach the microtubule end. The reduction for assisting loads is expected since they suppress backward motion and the microtubule end terminates forward-biased slip events. Surprisingly, even at assisting loads that were in magnitude significantly above the stall force of single motors, Kip3 still took single 8-nm center of mass steps against the force. Since such steps were not observed on the MT lattice, the observation suggests a change in the binding affinity at the microtubule end consistent with the switching model (21). The role of the microtubule end exploration is unclear, but might enable sideward stepping (17, 19) to find a protofilament that protrudes beyond the last helix of tubulin dimers that all have neighbors (Fig. 5). Since such a protruding protofilament lacks stabilization through lateral tubulin bonds it is curved and might represent the “weakest” protofilament most suitable for depolymerization. Even for blunt MTs, there is at least always one tubulin dimer at the MT seam that has only one lateral bond. We used GMPCPP-stabilized MTs that have mostly blunt, but flared ends with curved protofilaments similar to MTs grown in GTP (35, 36). Furthermore, independent of the nucleotide state, tubulin is thought to be in a bent state when it does not have neighbors bound (36). According to the switching model (21), the ATPase activity at the microtubule end on a curved protofilament is suppressed. If curved tubulins suppress ATPase activity, motors should not translocate beyond the first encountered curved tubulin dimer. Thus, Kip3 may not reach the end of a long curled protofilament. Instead, the front motor head should be in the ATP state and strongly bound to the first curved tubulin dimer. According to the consensus kinesin stepping model (37), the rear head should have initially ADP bound. Since hydrolysis in the front head is suppressed (21), ADP in the rear head might eventually be released leading to a rigor bound rear head. High assisting loads might induce stepping of the motor with both heads being in a strongly bound state (38). Another possible explanation for the steps at high forces, might be occasional ATP hydrolysis accompanied by nucleotide exchange.

**FIGURE 5.**
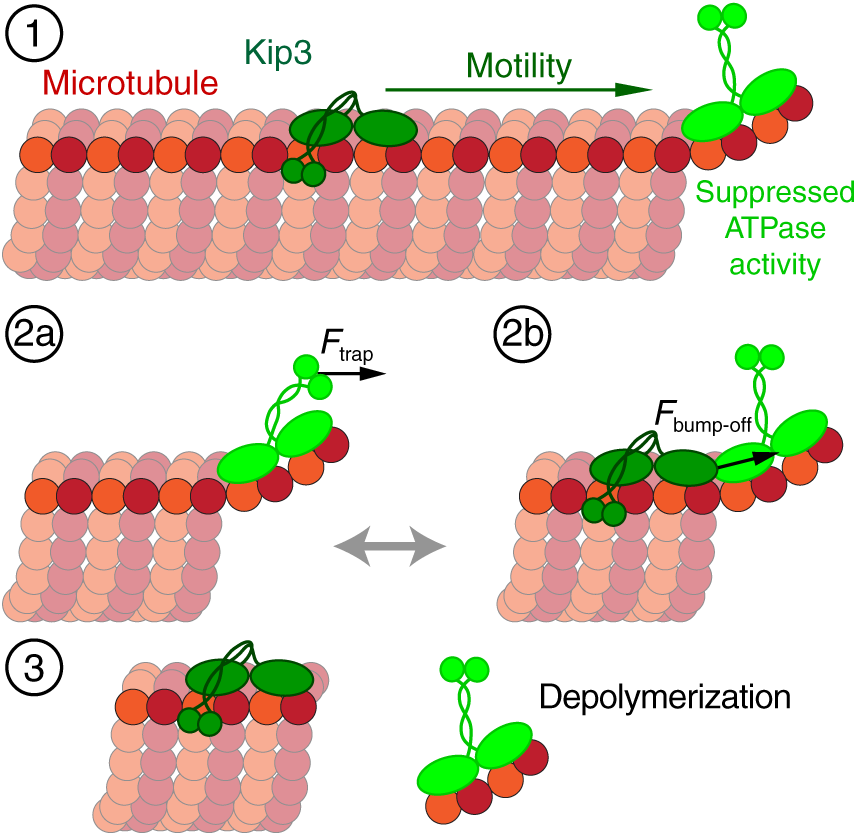
Model of collective, force-dependent microtubule depolymerization. (*State 1*) Kip3 (dark green) walks to the MT plus end with its processivity enhanced by the second microtubule binding domain in the tail and in contact with the microtubule lattice. At a curved single protofilament overhang, Kip3 switches to a strongly bound state with suppressed ATPase activity (bright green motor). A Kip3-tubulin complex may be removed either (*State 2a*) through an external force *F*_trap_, e.g. applied via an optical trap or equivalently (*State 2b*) through an incoming Kip3 that applies an internal “bump-off” force *F*_bump-off_. The second microtubule binding domain of the last motor on a protruding protofilament may not have access to the microtubule lattice or be “bumped-off” as well. (*State 3*) The strongly bound Kip3 was detached removing 2 tubulin dimers.

Loads decreased the end residence time of motors. For low assisting loads up to about +0.5 pN towards the microtubule plus end, the sensitivity to force was much larger compared to hindering loads of the same magnitude or higher assisting loads (Fig. 3A). This asymmetry may be due to the second microtubule binding site in the tail of Kip3. This site could still be in contact with the MT lattice for hindering and small assisting loads but not for high assisting loads. In the latter case, the Kip3 tail is most likely pulled away and beyond the end of the microtubule in our experiments. Because the end residence times were generally smaller for assisting loads, this binding site may contribute to the long Kip3 microtubule end dwelling relevant under physiological conditions and consistent with previous work (14). In support of this interpretation, the reported zero-load end residence time of Kip3 that lacks the tail microtubule binding site of 16.3 ± 0.6 s (14) is in excellent agreement with our high-assisting-load data extrapolated to a zero-load end residence time of 17 ± 2 s (Fig. 3A). Together with the extrapolation of the run length to the zero-force run length of the Kip3∆T-LZ mutant (Fig. 3B), these findings indirectly confirm the relevance of the tail MT-binding site for Kip3’s high processivity and end residence.

Is Kip3’s microtubule end activity different from the normal stepping mechanism? To answer this question, we compared the end residence time with run times on the microtubule lattice. The run times of Kip3 on the lattice were consistently higher than end residence times at the same loads with a similar distance-to-transition-state. Shorter end residence times imply a mechanism at the microtubule end that is different from the motility mechanism on the lattice. This finding is consistent with the switching mechanism (21)— depolymerization by Kip3 does not require motile Kip3 but is coupled to a switch in the binding state with suppressed ATPase activity. In our experiments, we cannot directly distinguish whether motors at the microtubule end simply detached from the microtubule or whether a motor-tubulin complex dissociated. Yet, we measured an increased depolymerization speed when MT ends were exposed to single Kip3 motors subjected to loads (Fig. 4). Thus, even though we cannot tell whether tubulin was removed at every end dissociation event, motors under load did significantly promote tubulin removal. Furthermore, based on the low off-rate of the GFP-nanobody interaction (33), we do not expect that the GFP-tagged motor dissociated from the GFP nanobody during microtubule end events. If we assume that motors bound to bent protofilaments have switched to an ATP-bound, but hydrolysis incompetent state as demonstrated in the switching model (21), motors should be strongly bound (Fig. 5). Since strongly bound motors on the lattice (AMPPNP data in Fig. 3A) took more than an order of magnitude longer to detach compared to motors at the end, we favor the hypothesis that motors dissociated from the end in a complex with tubulin and that force promoted the breaking-off of tubulin-tubulin bonds. How many tubulin dimers are removed per dissociation event is unclear. In the absence of ATP hydrolysis, we expect that eventually the second head dissociates its bound ADP. Then, both heads are in a strongly bound state and the dissociated complex should contain at least two dimers. Potentially, more dimers might be removed because motors, according to the switching model, will stop at the encounter of the first curved tubulin dimer. Thus, if the curled protofilament extends beyond that dimer all of those tubulins might be removed along with the motor. On the other hand, if many motors are present at the microtubule end, we do not expect many long protruding curved protofilaments. Also, there might not be sufficient time for ADP dissociation such that only one head is strongly bound and consequently only one dimer is removed per motor consistent with the reported 1–2 dimers/motor (13). A direct observation of tubulin removal by Kip3 remains an open experimental challenge. In a combined TIRF-optical tweezers setup (39), we tried to use MTs composed of 100 % fluorescently labeled tubulin and detect whether motors dissociated in a complex with tubulin. However, several factors precluded such an approach: motor motility was strongly reduced on such microtubules, autofluorescence of trapped microsopheres and bleaching in the optical tweezers limited the sensitivity, and, without trap, dissociated molecules rapidly diffused out of the TIRF field.

Since both load and increasing motor concentration accelerates microtubule end detachment, we can ask what load—in our case applied through the tail of the motor—on a single motor has the same effect as having many motors at the microtubule end that depolymerize the microtubule collectively. Therefore, we calculated the average time needed to remove a tubulin dimer. Based on the maximal Kip3-induced microtubule depolymerization speed of about 15 nm/s (13) or 8 nm/s for the mutant that lacks the second MT binding site (14), this time is ≈0.5–1 s (*τ*_end_ = 8 nm / depolymerization speed). Based on the exponential decay of the end residence time for assisting loads, this removal time corresponds to a tail-applied force of ≈4–5 pN, the lower force limit corresponding to the depolymerization rate without the second microtubule binding domain. At this force, the end residence time is about 80× shorter compared to the one in the absence of loads. This force range is also much higher than the stall force of a single Kip3 suggesting that a single motor cannot break-off tubulins. While multiple motors can generate higher forces (17), it is unclear how such an augmented force might be transmitted to the last motor and contribute to the depolymerization. Instead, we expect that a single motor should stabilize the last dimer through the additional connection via the motor’s neck linker to the previous tubulin dimer. If a second motor binds directly behind—either through its own motility or as in (21) as a monomer out of solution, steric hinderance—as suggested by structural modelling (18)—in particular, for a curved protofilament may generate sufficient force to bump-off the front motor-tubulin complex (Fig. 5). Since we applied forces through the tail, a direct comparison to forces that may arise from steric hindrance is difficult. Nevertheless, we measured the largest detachment sensitivity to force for assisting loads (Fig. 3A), i.e. a force directed towards the microtubule plus end. For the bump-off model, we expect that forces due to steric hindrance act in the same direction. One requirement for the bump-off is that the motor heads are in a strongly bound state. Otherwise, heads might be only “bumped off” of tubulin resulting in a motor detachment without tubulin removal. This requirement seems plausible based on the current kinesin stepping model (37) and the Kip3 switch mechanism (21). Thus, for effective tubulin removal both the bump-off and switching mechanisms are necessary. One additional aspect of the search for the “weakest” tubulin at the microtubule end and the bump-off model, in particular, at high concentrations that imply a kind of “traffic jam”, is that the second microtubule binding site of the last Kip3 might also be displaced from the microtubule, disconnecting the last, strongly bound Kip3 from the remaining microtubule lattice (State 2b in Fig. 5). Thus, our data provide evidence that both the switching and bump-off mechanism may contribute to microtubule depolymerization through ensuring a strongly-bound nucleotide state of the motor and an applied force through steric hindrance, respectively.

In summary, the force dependence of the end residence time suggests a combination of the bump-off model (13) with the switching model (21). Our combined model suggests that a single motor at the microtubule end—most likely having two strongly bound heads together with a tether to the remaining lattice via its second microtubule binding site—stabilizes microtubules. For efficient dissociation of a motor-tubulin complex, another motor is required to apply a force on the last motor. While the amount of force that is acting between an individual pair of consecutive motors will always be the same, the effective time-averaged force will depend on the motor concentration and, therefore, on the motor arrival rate at the end. In this sense, the concentration dependence of the end residence time is equivalent to its force dependence. Thus, collectively microtubules can be efficiently depolymerized in a force-dependent manner.

## SUPPORTING MATERIAL

Supporting material is available at http://www.biophysj.org/biophysj/supplemental/XXX

## AUTHOR CONTRIBUTIONS

M.B. and A.J. performed measurements and analyzed the data. M.C. expressed and purified the Kip3. T.J.J. programmed the force clamp control software. M.B., A.J., and E.S. wrote the manuscript.

## ACKNOWLEDGMENTS

We thank Ulrich Rothbauer (NMI, Reutlingen, Germany) for providing the anti-GFP nanobody. We thank Carolina Carrasco Pulido for comments on the manuscript. Mayank Chugh acknowledges financial support from International Max Planck Research School “From Molecules to Organisms”, Max Planck Institute for Developmental Biology, Tübingen. This work was supported by the German Research Foundation (DFG, JA 2589/1-1, CRC1011, project A04), the Institutional Strategy of the University of Tübingen (Deutsche Forschungsgemeinschaft, ZUK 63), Carl Zeiss Foundation (Forschungsstrukturprogramm 2017), and the University of Tübingen.

## Supporting Material

**TABLE S1.** Distances to the transition state x^‡^ (nm). Errors are standard errors.

|  |  |
| --- | --- |
| End residence time, hindering | $4.3 \pm 0.6$ |
| End residence time, assisting | $2.9 \pm 0.1$ |
| Run time in ATP, assisting | $3.3 \pm 0.4$ |
| Attachment time in AMP-PNP | $3.0 \pm 0.4$ |
| Attachment time in ADP | $6.4 \pm 0.5$ |
| Run length, assisting | $1.9 \pm 0.4$ |
| End range $\Delta$, assisting | $1.5 \pm 0.2$ |

**FIGURE S1.**
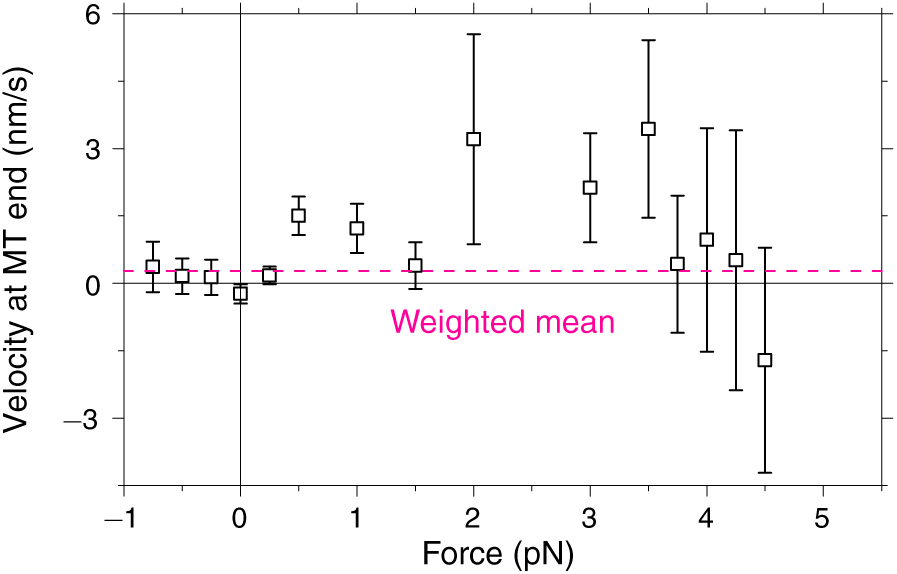
Kip3 velocity at the MT end as a function of load. Error bars are SEMs. The *magenta dashed line* refers to the overall weighted mean of 0.2 ± 0.1 nm/s.

**FIGURE S2.**
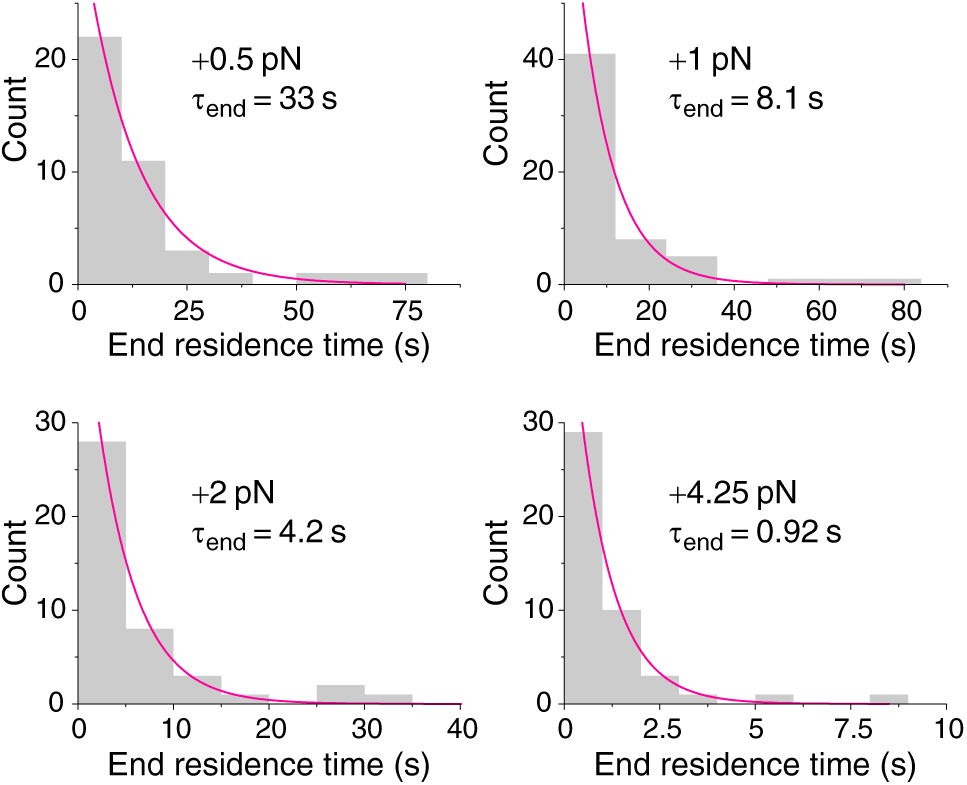
Distribution of end residence times as a function of assisting load. Four exemplary histograms of end residence times for different assisting loads with exponential fits and time constants *τ*_end_ corresponding to the mean values. Bartlett’s tests confirmed that end residence times were exponentially distributed.

**FIGURE S3.**
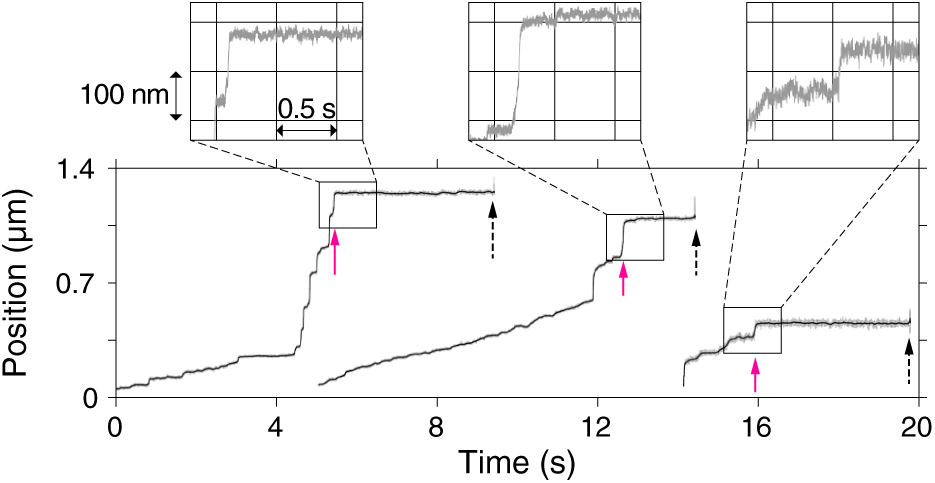
Kip3 slips to the microtubule plus end under assisting loads without instantaneous detachment. Three exemplary traces of Kip3 (+3.75 pN left and middle trace, +2 pN right trace, respectively), in which the motor slipped until it reached the end (*magenta arrows*). At the end, Kip3 did not detach immediately but after a prolonged end residence time (*black dashed arrows*). Insets show magnified slipping to the ends.

